# Primary cilia coordinate c-KIT signaling induced proliferation in alpha cells

**DOI:** 10.64898/2026.09.10.750659

**Authors:** Ganga Deshar, Mariam G Aslanyan, Gaurav D Diwan, Alexandros Karagiannopoulos, Tina Beyer, Fabiola Campestre, Alicia Vilas, Irene Cózar-Castellano, Lena Eliasson, Robert B Russell, Karsten Boldt, Ronald Roepman, Søren T Christensen, Lotte B Pedersen, Jakob G Knudsen

## Abstract

Circulating glucagon levels are elevated in patients with diabetes and obesity and contribute to hyperglycemia. The mechanisms underlying hyperglucagonemia remain poorly understood, but expansion of pancreatic α-cell mass is thought to play an important role. Primary cilia are sensory organelles that act as signaling hubs, for pathways controlling cell differentiation, proliferation, and function, yet their contribution to α-cell biology remains poorly defined. Here we investigated the role of primary cilia in the regulation of α-cell proliferation. To this end we generated the first α-cell primary cilia proteome identifying 167 cilia-enriched proteins. Among these, we identified and validated the proto-oncogene receptor tyrosine kinase, c-KIT, as a novel ciliary receptor in α-cells. We further show that c-KIT signaling depends on intact primary cilia and that stimulation of isolated mouse islets with its endogenous ligand, stem cell factor (SCF), promotes α-cell proliferation. In human pancreatic islets KITLG mRNA expression, but not KIT mRNA expression, correlated positively with donor BMI, suggesting that increased ligand availability drives c-KIT signaling in obesity. Together, our findings identify the primary cilium as a signaling platform c-KIT in α cells and reveal a ciliary axis that may drive α-cell expansion and hyperglucagonemia in obesity and diabetes.

## Introduction

Diabetes is characterized by elevated blood glucose levels and impaired regulation of insulin and glucagon secretion from pancreatic islets. Circulating glucagon levels are already elevated individuals with obesity and prediabetes (Starke et al., 1984, Færch et al., 2008) and is thought to contribute to the development of hyperglycemia. Although the mechanisms underlying hyperglucagonemia remain incompletely understood, defects in both intrinsic and paracrine control of pancreatic α-cells are thought to be central. While intrinsic metabolic dysfunction impairs glucagon secretion in diabetes (Dai et al., 2022, Knudsen et al., 2019, Zhang et al., 2013), somatostatin resistance (Kellard et al., 2020, Omar-Hmeadi et al., 2020) and increased α-cell proliferation (Ruiz-Pino et al., 2025) have been proposed to underlie the development of hyperglucagonemia. However, the molecular mechanisms linking obesity to altered paracrine signaling and α-cell mass expansion remain poorly defined.

Primary cilia are microtubule-based, antenna-like organelles that extend from the mother centriole at the surface and act as specialized sensory hubs integrating developmental and homeostatic pathways. They are present in a wide range of tissues, including metabolically active organs such as pancreatic islets and hypothalamus (Mill et al., 2023, Wachten and Christensen, 2025). Multiple signaling pathways are organized through primary cilia, including Hedgehog, G protein-coupled receptor (GPCR), receptor tyrosine kinase (RTK) and TGFβ superfamily signaling (Wachten and Christensen, 2025, Anvarian et al., 2019). Pathogenic variants in cilium-associated genes cause more than 35 pleiotropic human disorders, collectively known as ciliopathies, which frequently present in kidney, brain, and cardiac phenotypes, as well as metabolic dysfunction and obesity (Reiter and Leroux, 2017, Mill et al., 2023, Thomsen et al., 2025).

Most, if not all, pancreatic islet cell types possess primary cilia, which have emerged as important regulators of both insulin and glucagon secretion. Somatostatin receptor 3 (SSTR3), is a well-established ciliary GPCR (Berbari et al., 2008), enriched in the primary cilia of both α and β-cells (Iwanaga et al., 2011), where it mediates paracrine communication with δ-cells (Nilsson et al., 2025, Sanchez et al., 2022). Similarly, the GPCRs free fatty acid receptor (FFAR) 1 and FFAR4 amplify insulin and glucagon secretion, respectively, through ciliary signaling in intact islets (Wu et al., 2021). Beyond the β-cell cilium’s role as a Ca²⁺ compartment for paracrine GABA signaling (Sanchez et al., 2022), RTKs, including PDGFRα, IGF-1R and FGFR family members, also traffic to primary cilia in other cell types (Schneider et al., 2005; Yeh et al., 2013; Clement et al., 2013; Martin et al., 2018; Nita et al., 2025), raising the possibility that ciliary RTK signaling operates in islet endocrine cells as well.

Primary cilia are themselves highly dynamic organelles as their assembly and disassembly are tightly coupled to the cell cycle and developmental cues. In most vertebrate cells, the cilium is disassembled before mitosis as the centrosome is repurposed to organize the mitotic spindle (Mill et al., 2023, Ford et al., 2018, Thomsen et al., 2025). Because this cell-cycle-linked organelle also hosts RTK and GPCR signaling, we reasoned that primary cilia may regulate not only α-cell function but also α-cell proliferation. Here, we define the first α-cell primary cilia proteome and identify the proto-oncogene RTK c-KIT as a novel ciliary receptor in α cells. We show that c-KIT/ERK signaling and α-cell proliferation require intact primary cilia, and we provide evidence that ciliary c-KIT may contribute to α-cell expansion in obesity and type 2 diabetes.

## Materials and Methods

### Animal experiments

All animal experiments were approved by the Danish Animal Inspectorate. Female WT C57B6Nrj mice were used in all experiments. The animals were housed in a 12 hour-12 hour light-dark cycle with ad libitum access to standard rodent chow and water. All animals were used between the age of 12 and 20 weeks.

### Cell culture

Primary mouse embryonic fibroblasts (MEFs) were cultured in DMEM and 45% F12 L-glutamine (InVitrogen, Taastrup, Denmark) supplemented with 10% heat-inactivated fetal calf serum (FCS) and 1% penicillin-streptomycin at 37 °C, 5% CO_2_. The αTC1-6/9 cell lines (αTC1-6; CRL-2934, ATCC), were cultured in RPMI 1640 (118279-020, Gibco, Thermo Fisher Scientific) containing 15 mM Hepes (15630-56, Gibco, Thermo Fisher Scientific), 1% penicillin/streptomycin (15140-122, Gibco, Thermo Fisher Scientific), 10% fetal bovine serum (FBS; 10270-10, Gibco, Thermo Fisher Scientific) and 11 mM glucose at 37 °C with 5% CO_2_.

### Cell line generation

Lentiviral particles carrying pLenti-ARL13B-TurboID-3xFlag or pLenti-miniTurboID were generated in HEK293T cells as described previously (Rezi et al., 2024), and RPMI 1640 culture medium was used to transduce αTC1-6 cells, followed by selection with Blasticidin, to generate αTC1-6 cell lines stably expressing ARL13B-TurboID-3xFlag or miniTurboID.

### Proximity labelling

Proximity labelling was performed in the ARL13B-TurboID-3xFlag αTC1-6 cell line, with the matched miniTurboID αTC1-6 cell line as negative control, in six independent biological replicates. Cells were seeded at ∼15% confluency and cultured for 24 h in normal RPMI 1640 medium supplemented as described above, then switched to ciliogenesis-inducing starvation medium (RPMI 1640 with 0.2% FCS) for 48 h. 10uM biotin was added overnight to induce biotinylation.

(Sigma-Aldrich, B4501).

After labelling, cells were washed three times in ice-cold PBS to remove free biotin and lysed in RIPA buffer (50 mM Tris-HCl pH 7.5, 150 mM NaCl, 1% NP-40, 0.1% SDS (Life Technologies, 15553-027), 0.5% Sodium Deoxycholate, 1 mM UltraPure EDTA pH 8.0 (Invitrogen, 11568896)), followed by sonication and 1 h of rotation at 4 °C. Lysates were cleared by centrifugation at 14,000 rpm for 20 min at 4 °C, snap frozen and stored at −80°C until sample enrichment. Biotinylated proteins were enriched for 2 h at 4 °C on StrepTactin Superflow beads (IBA, 2-1206-025). Beads were washed twice in washing buffer (1x TBS (20 mM Tris, 150 mM NaCl) containing 0.1% NP-40) at 4 °C followed by two washes in 1xTBS at 4 °C. On-bead tryptic digestion was carried out in a Thermomoxer for 1 h at 27 °C, with agitation at 800 rpm, in Trypsin digestion buffer (2 M Urea, 50 mM Tris-HCl (pH 7.5), 5 μg/sample trypsin (Serva, 9002-07-7)). The samples were digested further overnight (2 M urea, 50 mM Tris-HCl pH 7.5 and 1 mM DTT) at 22°C. The eluted peptides were snap frozen and stored at -80 °C until downstream mass spectrometry (MS) analysis.

### Mass spectrometry analysis and statistical identification of enriched proteins

The samples were analyzed and proteins were identified according to the method described in (Aslanyan et al., 2023, Whiting et al., 2026). Briefly, tryptic peptides were processed for LC-MS/MS using data-independent acquisition mode (DIA) on the QExactive Orbitrap mass spectrometer (Thermo Fisher Scientific). Raw data files were analyzed using DIA-NN 1.9 (Demichev et al., 2020) against the mouse database (Swiss-Prot, May 2024, #17,195 proteins), and high-accuracy spectra with a minimum false discovery rate (FDR) of 0.01 and tryptic peptides were used for protein abundance label free quantification. For proteomics data analysis, we used a custom in-house R script that replicates the analysis using the Perseus software (Tyanova et al., 2016). The LFQ intensity values were compared for miniTurbo ID samples versus those for ARL13b-TurboID samples. For samples where LFQ intensity values were zero in less than half of the replicates, while having nonzero LFQ intensity values in the other replicates, imputed values were generated drawn from a normal distribution that had a mean that was 1.8 times below the mean of the nonzero values and a standard deviation that was 0.5 times the mean. Subsequently, Student’s *t*-test was used for statistical comparisons between the LFQ intensity values of samples as well as the significance A test to infer samples with outlier log2 ratios (high or low). After removing the proteins that were significantly altered in the miniTurboID comparison, we devised a three-tier system to classify significant proteins from the ARL13b-TurboID comparison. Tier 1 proteins were ones where the corrected *P*-values (Benjamini–Hochberg correction) from the *t*-test were <0.05 as well as significance A test *P*-values were <0.05. Tier 2 proteins included proteins that only had significance A test *P*-values < 0.05 and Tier 3 proteins were the ones that only had corrected *P*-values from the *t*-test <0.05. For the dataset comparison with published ciliary proteomes, we downloaded the relevant supplementary files and tables to annotate the proteins in Tier 1 and Tier 2 from this study. For the Breslow et al. study, we obtained the mouse orthologs (Diwan et al., 2026) of the human proteins.

### STRING network analysis

STRING network analysis (https://string-db.org/) was used to identify functional protein association networks enriched in the ciliary fraction from the proximity labeling experiments. Briefly, the STRING database was searched using the 167 cilia enriched proteins from Tier 1 and 2. STRING carries out enrichment against the GO database, and we focused on the output of the GO biological process analysis. The results were ordered by Strength which is defined as log10(observed/expected) of proteins annotated by a term, thereby measuring how large the enrichment effect is. Terms were not grouped by similarity and overlapping members from the “Cell division” term was displayed in an interaction network.

### Islet isolation

Mice were euthanized by cervical dislocation, and islets were isolated by injection of 1.5 ml Liberase (1 mg/ml) into the common bile duct as previously described (Briant et al., 2018). Injected pancreata were digested at 37 °C for 12 min and subjected to mechanical disruption. Islets were handpicked in 7 mM glucose RPMI 1640 (Gibco, 61870-010) 10% FBS, 1% penicillin/streptomycin) and incubated at 37 °C in 5% CO_2_ for 1 h before initiation of experiments.

### Immunofluorescence microscopy analysis

αTC1-6 cells were cultured in RPMI 1640 as described above and seeded on coverslips; to induce cilia formation cells were serum-starved in 1% FBS for 48 h. Isolated islets or cells were washed in PBS and fixed in 4% PFA for 15 min followed by extensive washing in PBS. After permeabilization with 0.1% Triton X-100 and blocking with 5% Bovine Serum Albumin (BSA), cells were incubated in primary antibodies at 4 °C overnight. The primary antibodies used recognize c-KIT (Proteintech, 18696-1-AP), Glucagon (Sigma, G2654), ARL13B (Proteintech, 30332-1-AP), Acetylated α-tubulin (Merck, #T7451) and Ki67 (Cell signaling, #9129). Following this, cells were washed in PBS with 0.05% Tween 20 (PBST) and incubated with secondary Alexa fluor antibodies and washed in PBST. The cells were then mounted using Vectashield. Images were acquired using a Nikon Ti2 CrestV3 spinning Disc confocal microscope at 40x magnification. Image analysis was performed in ImageJ.

### Immunoblotting

αTC1-6 cells or primary MEFs were cultured as above and serum starved in 1% FBS for 48 h before the experiment. The media was removed and cells were washed gently two times with PBS at room temperature (RT) then incubated in media (1%FBS) supplemented with Stem Cell Factor (SCF) 50ng/mL for 3, 10 or 30 min at 37 °C and 10% CO_2_. Cells were then lysed in lysis buffer (150 mM NaCl, 20 mM Hepes, 1 mM EDTA, 10% glycerol, 0.5% Triton X-100, 1% SDS, protease and phosphatase inhibitor cocktail (Thermo Fisher Scientific, Cat. No. 78440)), sonicated 3 times for 10 sec each, and centrifuged at 15,000 g for 15 min. Cell lysates were collected and stored at -80 °C. Protein concentrations were measured using a Bradford Assay (Bio-Rad, DC protein assay Reagent B, # 5000114 DC Protein Assay Reagent A # 5000113, DC Protein Assay Reagent S # 5000115). Samples were prepared by adding 50 mM dithiothreitol (DTT), sample buffer (NuPAGE, Cat. No. NP0007, Invitrogen) and heated at 95 °C for 5 minutes. Protein samples were loaded in polyacrylamide gels (NuPAGE 4-10% Bis-Tris 12 well, Thermo Fisher Scientific Cat. No. NP0322box) and then transferred to a PVDF membrane (Invitrogen, Thermo Fisher Scientific Cat. No. LC2005). Membranes were stained with Ponceau and blocked in Tris-buffered saline with 0.1% Tween 20 (TBST) buffer with 5% skimmed milk for 1 h at 37^0^C on a shaker. Primary antibodies against ERK1/2 (Cell Signaling, 9102), ERK1/2-p (Cell Signaling, 9101) or DNTC (BD Biosciences, 610474), were added overnight at 4 °C. The next day, membranes were washed three times with TBST for 5 min each, followed by incubation with appropriate HRP conjugated secondary antibodies. Bands were visualized with Fusion FX Spectra (Vilber Lourmat, Eberhardzell, DE).

### Ciliation and ciliary length analysis

Cilia frequency and length were analysed using FIJI ImageJ software. Frequency was quantified by dividing the number of cilia per field of view by the number of nuclei (DAPI). Cilia and nuclei were manually counted using the “Multi-point” tool in ImageJ. Ten immunofluorescence microscopy (IFM) images were analysed per condition per replicate (n = 3). Cilia length was manually measured by tracing a line along the midline of each cilium using the “Segmented Line” tool in ImageJ. Thirty cilia were analysed per condition per replicate (n = 2).

### Hormone Secretion

Glucagon secretion was measured from groups of 10 islets/replicate. The islets were washed once with Krebs ringer buffer (KRB) (140mM NaCl, 3.6mM KCl, 1.3mM CaCl_2_, 0.5mM MgSO_4_, 10mM HEPES, 0.5mM NaH_2_PO_4_, and 25mM NaHCO_3_ adjusted pH 7.4, with 0.36 mmol/L NEFA bound to 6.6% fatty acids free BSA (Sigma)) with 5 mmol/L glucose KRB, for 1 h at 37°C in 5% CO_2_. Islets were then incubated sequentially at 1 mmol/L and 10 mmol/L glucose with or without 50ng/ml SCF. The supernatant was collected, and islets were harvested in acid ethanol and sonicated. Glucagon concentrations were determined using a Homogeneous Time-Resolved Fluorescence kit detecting glucagon (Revvity; 62CGLPEG), according to the manufacturer’s instructions.

### RNA-seq dataset and Differential expression analyses

Normalized gene expression data and donor phenotype information were obtained from a publicly available bulk RNA-sequencing dataset of 219 human pancreatic islet donors (Bacos et al., 2023). All gene expression analyses were performed using DESeq2 normalized counts.

Donors were classified into three glycemic groups based on HbA1c levels and T2D diagnosis: normoglycemic (ND; no T2D diagnosis and HbA1c < 42 mmol/mol, n = 149), impaired glucose tolerance (IGT; no T2D diagnosis and 42 ≤ HbA1c < 48 mmol/mol, n = 37), and type 2 diabetes (T2D; confirmed T2D diagnosis, n = 33). Non-diabetic donors with missing HbA1c values were assigned to the ND group. For three-group comparisons (ND vs IGT vs T2D), all pairwise unpaired Mann-Whitney U tests were performed. Donors were also stratified by BMI into five categories (< 20, n = 5; [20, 25), n = 79; [25, 30), n = 97; [30, 35), n = 28; > 35 kg/m², n = 9). One donor with a missing BMI record was excluded. Differences in expression across BMI groups were evaluated with a global Kruskal-Wallis test, followed by pairwise unpaired Mann-Whitney U post-hoc tests.

### Correlation analyses

To assess relationships between target gene expression and continuous variables or co-expressed genes, pairwise Pearson correlations were calculated by using DESeq2 counts normalized by variance-stabilizing transformation (VST). Scatter plots were generated with linear regression overlays and data points are color-coded by glycemic group (ND, IGT, T2D) using the R package ggpubr (v0.6.2).

### Statistical analysis

All statistics were carried out using GraphPad Prism 11 software (San Diego, CA) or R. Where appropriate, t-tests, one-way ANOVA and two-way ANOVA analysis were performed with post-hoc analysis to identify differences between groups.

## Results

### Profiling the primary cilium proteome of α-cells

Primary cilia are highly heterogeneous organelles whose proteomic composition varies considerably between cell types (Hansen et al., 2025). To investigate how primary cilia regulate α-cell biology, we applied unbiased cilium-targeted proximity labeling proteomics to define the α-cell ciliary proteome (Schermer et al., 2025). We generated αTC1-6 cell lines stably expressing TurboID fused to the ciliary membrane-associated protein ARL13B (ARL13B-TurboID) or miniTurboID alone (Figure S1A). Immunofluorescence microscopy (IFM) of serum-starved cells (to induce growth arrest and formation of primary cilia) incubated with biotin and stained for ARL13B together with fluorescent streptavidin demonstrated robust ciliary biotinylation in ARL13B-TurboID expressing cells (Figure S1B). Following serum starvation, biotin labeling, streptavidin pulldown, and mass spectrometry, we identified 3,516 proteins across six technical replicates. Of these, 167 were significantly enriched more than threefold in the ciliary fraction (Tier 1: P<0.05; Tier 2: SigA <0.05) (Figure 1A). The enriched proteins included well-established ciliary proteins like IFT140 (Cole et al., 1998) as well as transition zone and centrosomal components such as NPHP4 and CEP131 (Habbig et al., 2011, Andersen et al., 2003), validating the specificity of the approach. Comparison with previously published ciliary proteomes identified 77 proteins unique to the αTC1-6 ciliary data set (Figure 1B), supporting the existence of cell-specific ciliary signaling machineries. String network and Gene Ontology (GO) analyses of the 167 from Tier 1 and 2 revealed significant enrichment of proteins involved in cytoskeletal arrangement, cellular organization, development, and cell division (Figure 1 C). Among proteins associated with cell division (Figure 1D), we focused on the class III receptor tyrosine kinase Kit (c-KIT/CD117), a proto-oncogene with established roles in stem cell maintenance, proliferation, development and tumor progression (Lennartsson and Rönnstrand, 2012). Because c-KIT signaling has previously been implicated in β-cell proliferation (Feng et al., 2015), we hypothesized that ciliary c-KIT similarly could regulate α-cell proliferation.

**Figure 1.**
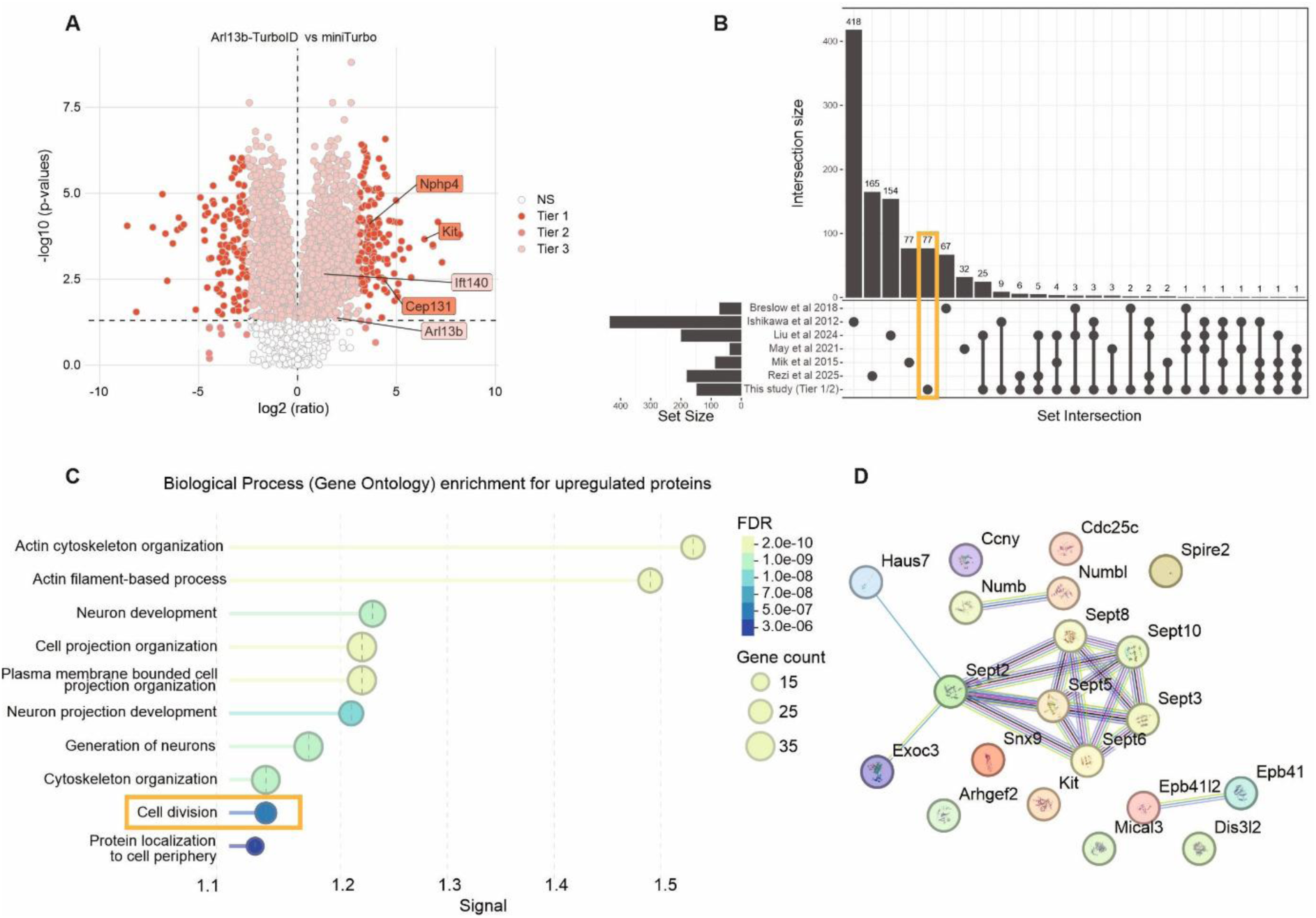
Profiling primary cilia protein expression in alpha cells. A) Volcano plot visualizing differential enrichment in primary cilia (positive Log2 ratio) and Cytoplasm (negative Log2 ratio). The proteins are colored according to their significance tier (Tier 1, 2, 3, and non-significant (NS)). (B) Upset plot comparing the obtained proteome with other with primary cilia proteomes to identify novel ciliary proteins. (C) Gene Ontology enrichment analysis for biological process performed using upregulated proteins in Tier 1 and 2. Figure shows GO terms listed according to FDR. (D) Protein interaction network analysis showing enriched proteins belonging to the GO term Cell division colored lines indicates origin of evidence for interaction prediction. known Interactions: curated database (Cyan), experimentally determined (magenta). Predicted interactions: Gene Neighborhood (green), gene fusions (red), gene co-concurrence (blue). Others: Text mining (light green), Co-expression (black); Protein homology (purple).

### c-KIT localizes to the primary cilium

To validate the ciliary localization of c-KIT, we first examined its expression in serum-starved αTC1-6 cells. Immunoblotting readily detected c-KIT protein (Figure 2A), while IFM demonstrated localization of c-KIT at the primary cilium and throughout the cell body (Figure 2B). Similarly, ciliary localization of c-KIT was observed in isolated mouse islets (Figure 2C). Co-staining for glucagon revealed that 98.5% of the c-KIT-positive cells in the isolated whole islets were α-cells (Figure 2D), with a small fraction of non-α-cells also expressing c-KIT, consistent with previous reports (Yashpal et al., 2004). Notably, most (73% ± 3.6%) α-cells expressed detectable c-KIT (Figure 2E), suggesting the existence of functionally distinct α-cell subpopulations as previously reported (Huang et al., 2026, Frueh et al., 2026).

**Figure 2.**
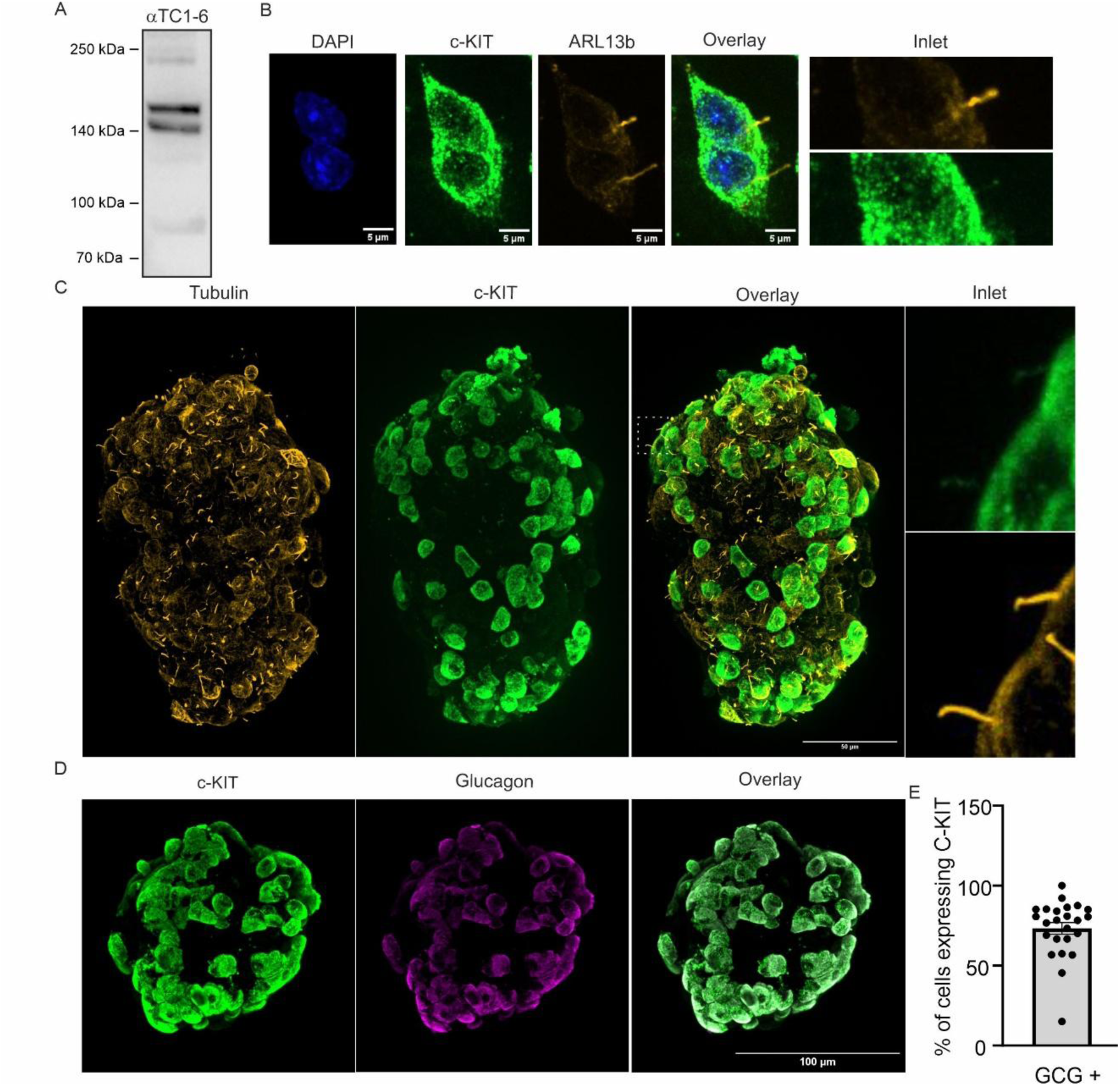
The RTK c-KIT localizes to the primary cilium. (A) Representative western blot of c-KIT and (B) immunofluorescent staining of c-KIT and ARL13B in ciliated αTC1-6 cells (scale bar is 5µm). Immunofluorescent staining of (C) acetylated α-tubulin and c-KIT (scale bar is 50µm), or (D) c-KIT and glucagon in isolated mouse islets (scale bar is 100µm). (E) Quantification of c-KIT expressing glucagon positive cells from (D) (n= 22 islets from 5 mice). All Data are presented as mean ± SEM. Scalebars are 10 µm.

### Activation of c-KIT signaling depends on the primary cilium

We next investigated whether stimulation with the endogenous c-KIT ligand Stem Cell Factor (SCF) activates c-KIT signaling in αTC1-6 cells. SCF induced rapid phosphorylation of the MAP kinases ERK1/2 in αTC1-6 cells, with robust activation after 3 and 10 min of stimulation (Figure 3A-C). Since ERK1/2 activation has previously been associated with ciliary disassembly (Dougherty et al., 2023, Ritter et al., 2019, Wang et al., 2013) we asked whether SCF signaling affected ciliary dynamics. SCF Treatment of αTC1-6 cells for 6 h and 10 h, led to significantly shorter cilia at 6 and 10h, without altering the proportion of ciliated cells (Figure 3D-F). To determine whether loss of primary cilia can induce proliferation in α-cells, we depleted the ciliary transport protein IFT88, which is essential for ciliogenesis (Schneider et al., 2005, Pazour et al., 2000, Casanueva-Álvarez et al., 2026), in αTC1-9 cells. Knockdown of IFT88 reduced ciliation frequency (Figure 3G and H) and increased cell proliferation (Figure 3I). To examine whether cilia are required for c-KIT signaling, we stimulated serum-starved wild-type and *Tg737*^orpk^ mutant MEFs with SCF. *Tg737*^orpk^ mutant MEFs express a hypomorphic IFT88 allele that results in absent or severely shortened primary cilia (Schneider et al., 2005). In wild-type MEFs, SFC induced rapid phosphorylation of ERK1/2 after 3 and 10 min (Figure 3J-L), while this response was abolished in *Tg737*^orpk^ MEFs (Figure 3 J-L). Interestingly, expression of total ERK1 (but not ERK2) was lower in Tg737orpk compared to WT MEFs (Figure 3J). Together these findings suggest that intact primary cilia are required for efficient c-KIT-mediated ERK1/2 activation.

**Figure 3.**
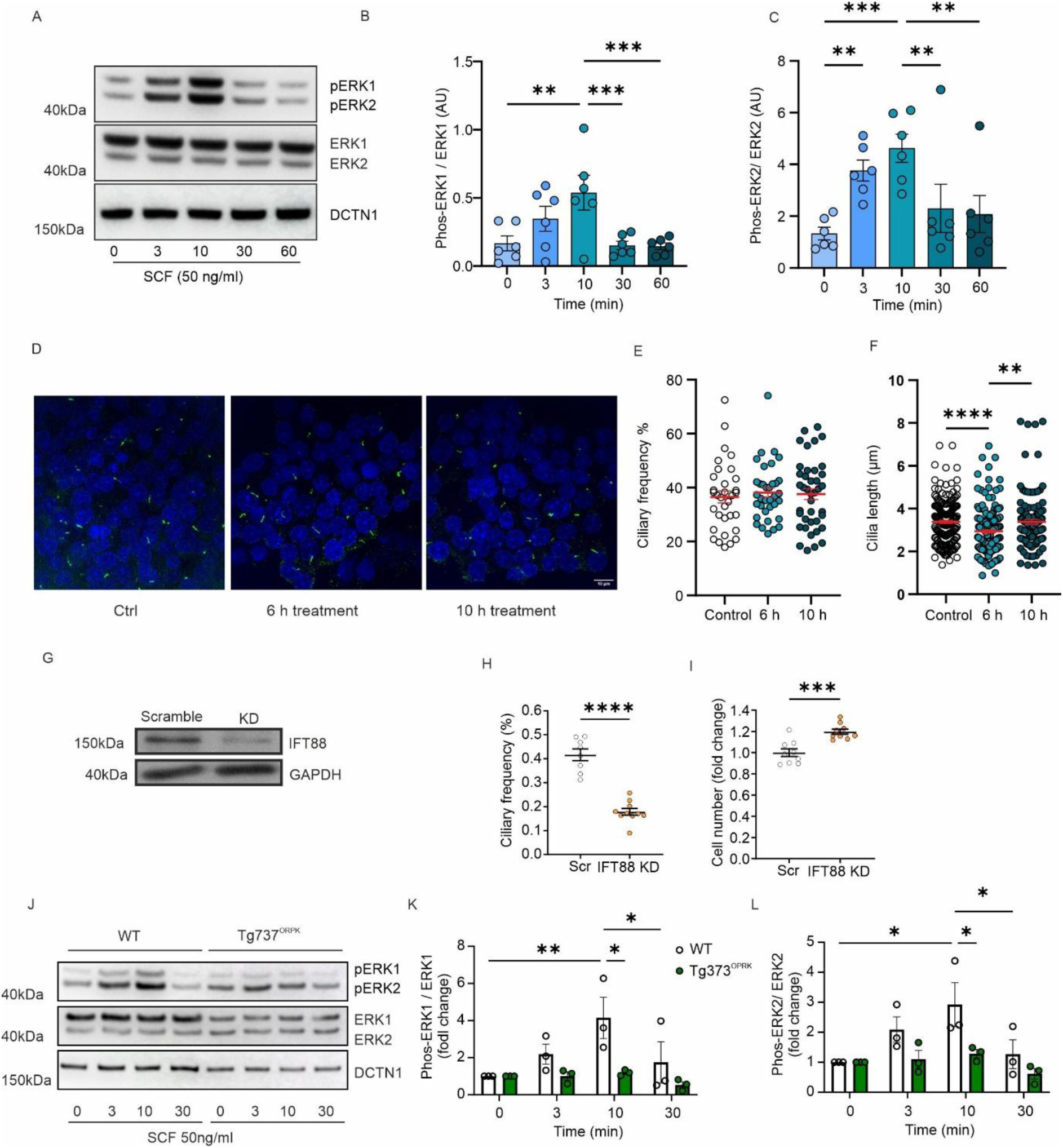
Activation of c-KIT signaling requires primary cilia localization. (A) Representative western blot analysis of ERK1/2 signaling in αTC1-6 cell after stimulation with SCF (50 ng/ml). (B, C) Quantitative analysis of data from (A) (n = 3 experiments from 3 different passages). (D) Immunofluorescent stanning of primary cilia with ARL13B (green), DAPI (blue) in 48 h serum-starved αTC1-6 cells treated with SCF (50 ng/ml) for 6 and 10 h. (E) Accumulated ciliary frequencies and (F) ciliary lengths from (D) (n= 33-40 fields of view for frequency, or n=120-180 cilia for lengths from 3 independent experiments). (G) Representative western blot of IFT88 knockdown in αTC1-9 cells. (H) Ciliary frequency in control (scramble) and IFT88 knockdown αTC1-9 cells (n = 7-10 replicates from 3 independent experiments). (I) Increase in cell number as a measure of proliferation in control (scramble) and IFT88 KD αTC1-9 cells (n=9 replicates from 3 independent experiments). (J) Representative western blot analysis of ERK1/2 signaling in wild type (WT) or *Tg737*^orpk^ MEFs after stimulation with SCF (50 ng/ml). (K, L) Quantitative analysis of data from (J) (n = 3 experiments from 3 independent experiments). All quantitative data are presented as mean ± SEM, * (P<0.05), ** (P<0.01), *** (P<0.001) ****(P<0.0001).

### SCF stimulates alpha cell proliferation

Because ERK1/2 is a central regulator of cell proliferation (Lavoie et al., 2020), we next examined whether SCF promotes α-cell proliferation. Whole isolated mouse islets were stimulated with SCF for 24 h and 72 h, resulting in a significant increase in the number of Ki-67-positive α-cells compared with untreated controls (Figure 4A-B, C; Figure S2A). In contrast, SCF did not affect Ki-67 expression in non-α-cells (Figure S2B), differing from previous reports (Feng et al., 2015). Acute SCF stimulation had no effect on glucagon secretion at 1 or 10 mM glucose (Figure 4C). These findings suggest that ciliary c-KIT signaling selectively promotes α-cell proliferation without acutely affecting glucagon secretion.

**Figure 4.**
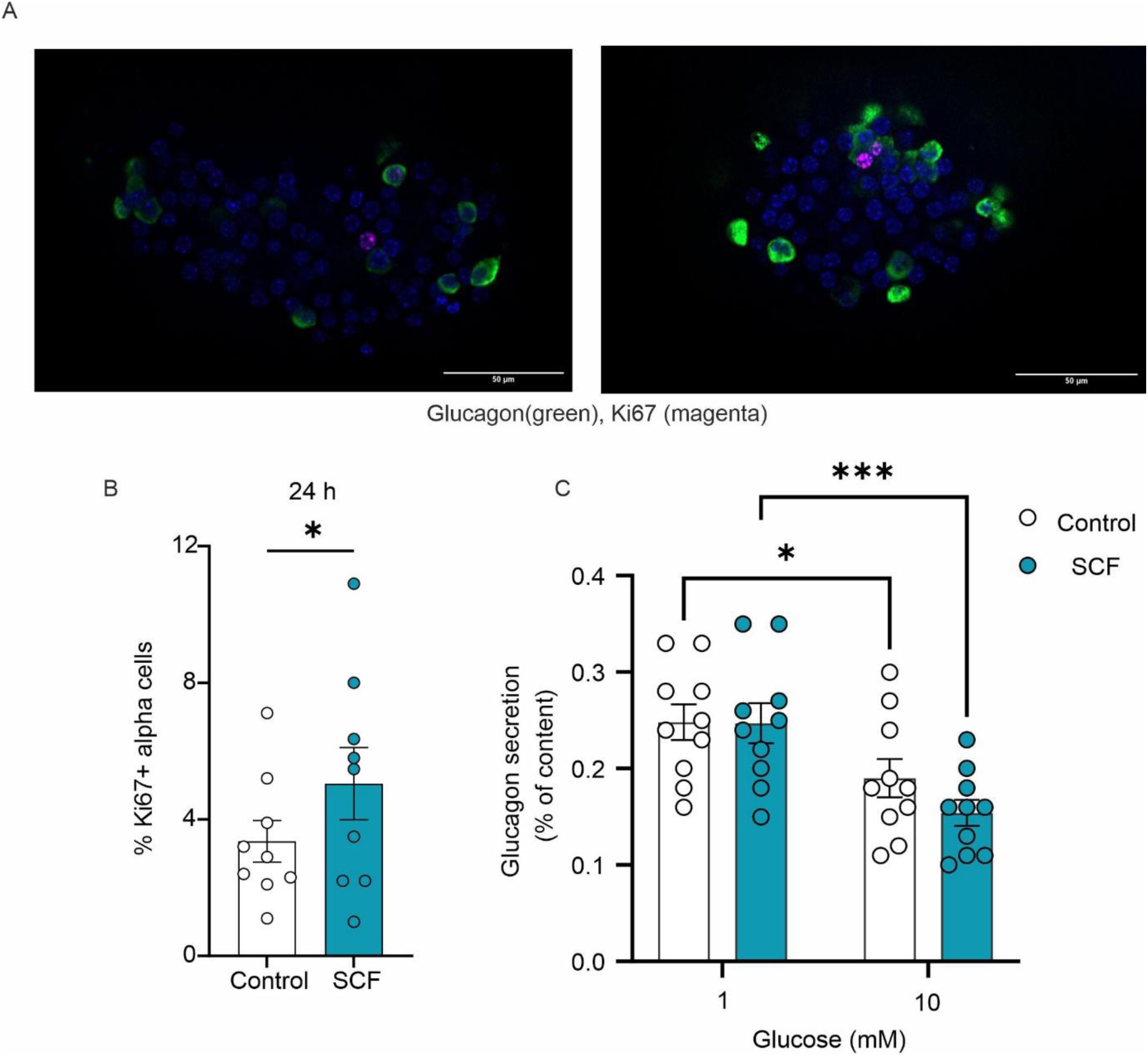
c-KIT stimulates alpha cell proliferation in whole islets. (A) Immunofluorescent staining of Ki67 (magenta), glucagon (green) and DAPI (blue) in isolated islets incubated in (left) vehicle or (right) SCF (50 ng/ml) for 24 h (scale bar is 50µm). (B) Accumulated data from (A) (n=9 mice from 3 separate experiments). (C) Glucagon secretion from isolated mouse islets in response to acute stimulation with SCF (50 ng/ml) at 1 and 10 mM glucose (n=10 mice from 4 separate experiments). All data are presented as mean ± SEM * (P<0.05), ** (P<0.01), *** (P<0.001).

### Stem cell factor (*KITL*) expression correlates with BMI

To explore the potential relevance of c-KIT signaling in human obesity and diabetes, we analyzed published islet RNA-sequencing datasets from human donors with impaired glucose tolerance (IGT) and type 2 diabetes (T2D) (Bacos et al., 2023). Expression of *KIT*, encoding c-KIT, did not correlate with body mass index (BMI), glycated hemoglobin (HbA1c) or diabetes status (Figure S3A-D), suggesting that altered receptor expression is unlikely to explain the changes in α-cell mass with obesity. SCF is encoded by *KITLG* and is primarily expressed by endothelial and perivascular cells (Ding et al., 2012). Because obesity is associated with increased islet vessel diameter (Dai et al., 2013) and thus higher numbers of endothelial cells, we examined whether *KITLG* expression correlated with metabolic parameters. Although *KITLG* expression did not correlate with HBA1c (Figure S3E-F), it showed strong positive correlation with BMI (Figure 5A). Stratification of donors into 5-unit BMI intervals, revealed higher *KITL* expression in individuals with a BMI between 30 and 35 compared to those with BMIs of 20-25 or 25-30, whereas individuals with a BMI below 20 exhibited markedly reduced intra-islet expression of *KITLG* (Figure 5B). These findings suggest that obesity is associated with elevated intra-islet SCF levels and support the hypothesis that enhanced SCF/c-KIT signaling contributes to α-cell expansion in obesity.

**Figure 5.**
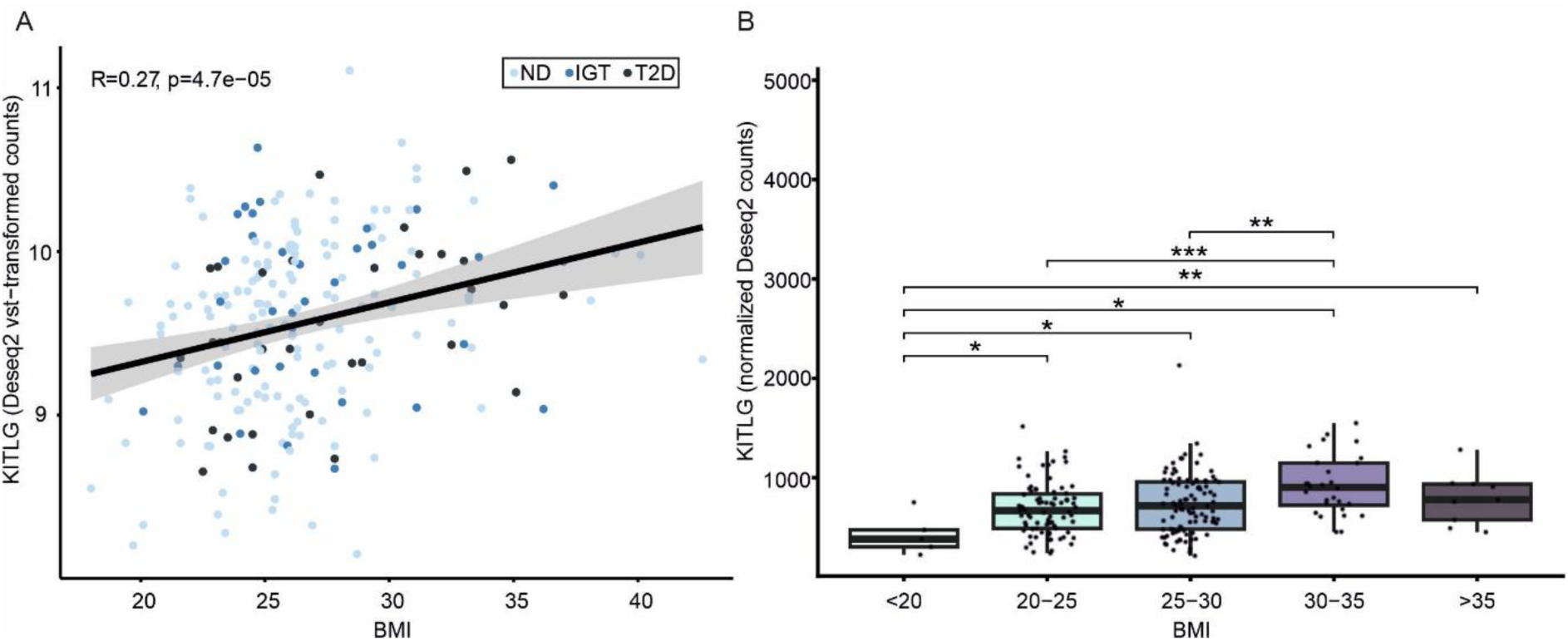
Stem cell factor expression in human islets correlates with BMI. (A) Correlation between expression of SCF in whole human islets and body mass index (BMI) in non-diabetic subjects (ND, n=149), subjects with impaired glucose tolerance (IGT, n=37) or Type 2 diabetes (T2D, n=33). (B) Data form (A) stratified for BMI (< 20, n = 5; [20, 25), n = 79; [25, 30), n = 97; [30, 35), n = 28; > 35, n = 9). Boxplots represent the median, the interquartile range (IQR; 25^th^ to 75^th^ percentiles) and whisker extending to the data range within 1.5 x IQR. * (P<0.05), ** (P<0.01), *** (P<0.001).

## Discussion

In this study, we identify c-KIT as a ciliary receptor tyrosine kinase in pancreatic α cells and show that SCF signaling through the ciliary c-KIT axis promotes ciliary shortening and α-cell proliferation. Using cilium-targeted proximity-labeling proteomics in αTC1-6 cells, we mapped 167 ciliary proteins including the RTK c-KIT. c-KIT localized to primary cilia in mouse islets and activation of c-KIT promotes cilia shortening in cultured cells, while SCF-induced ERK1/2 activation was abolished in cilium-deficient cells. Consistent with a mitogenic ciliary c-KIT axis, SCF stimulated α-cell proliferation in intact mouse islets, and in islets from human donors, intra-islet *KITLG* expression correlated positively with BMI. Together, these results position SCF/c-KIT as a candidate paracrine driver of α-cell mass expansion in obesity and hyperglucagonemia.

Several receptor tyrosine kinases converge on the primary cilium. PDGFRα, insulin-like IGF1R, selected FGFRs, and the angiopoietin receptors TIE1/2, localize to primary cilia in a variety of cell types (Schneider et al., 2005, Neugebauer et al., 2009, Martin et al., 2018, Nita et al., 2025, Teilmann and Christensen, 2005). Our findings extend this repertoire by identifying c-KIT as an additional ciliary RTK in alpha cells. Interestingly, c-KIT appears to regulate ciliary dynamics differently from PDGFRα, despite both belonging to the class III RTK family. Whereas PDGFRα signaling has primarily been implicated in directional cell migration (Clement et al., 2013, Schneider et al., 2010), c-KIT activation induced ciliary shortening and stimulated α-cell proliferation. This phenotype resembles the effects of IGF1R signaling, which likewise promotes ciliary resorption during mitogenic stimulation (Yeh et al., 2013), suggesting that distinct ciliary RTKs couple extracellular cues to specific cellular responses. ERK1/2 signaling is a well-established regulator of ciliary disassembly through phosphorylation of proteins that control IFT and axonemal microtubule dynamics (Broekhuis et al., 2013, Chaya et al., 2025). It is therefore plausible that c-KIT promotes ciliary shortening through ERK1/2-dependent mechanisms, thereby coupling ciliary remodeling to cell-cycle entry and proliferation, although this hypothesis requires direct experimental validation.

In pancreatic islet cells, primary cilia-mediated signaling has primarily been linked to the regulation of hormone secretion, cell polarity and gene expression. This has been studied predominantly in β-cells, where primary cilia mediate somatostatin signaling through SSTR3 and contribute to GABA responses (Sanchez et al., 2022). In contrast, only FFAR4 (Wu et al., 2021), and IDE (Casanueva-Álvarez et al., 2026, Merino et al., 2022) have so far been implicated in ciliary signaling in α-cells in the regulation of glucagon secretion. Our findings therefore further support that α-cells possess specialized signaling mechanisms that distinguish them from other endocrine cell types. Together with previous observations that α-cells preferentially utilize aerobic glycolysis, rely on fatty acid oxidation to sustain ATP production, and exhibit high glutamine metabolism (Armour et al., 2023a, Briant et al., 2018, Knuth et al., 2024, Armour et al., 2023b, Dean, 2020), the prominent expression of c-KIT may represent another feature contributing to the remarkable proliferative capacity of α-cells that becomes evident during diabetes and obesity.

Expansion of α-cell mass has previously been linked to elevated circulating amino acids, particularly glutamine, resulting from impaired hepatic glucagon signaling and disruption of the liver–α-cell axis (Dean et al., 2017, Riahi et al., 2023, Kjeldsen et al., 2024, Wewer Albrechtsen et al., 2018). Our findings here suggest that regulation of α-cell proliferation is likely to be more complex. Under the experimental conditions used here, SCF-mediated stimulation of c-KIT promoted proliferation in α-cells within intact isolated islets. Together with recent evidence that amino acids alone are insufficient to directly stimulate α-cell proliferation in diabetic animals (Riahi et al., 2023), our results suggest that α-cell expansion in obesity and diabetes may require the integration of multiple signaling pathways. Consistent with this idea, intra-islet expression of SCF increased with higher BMI, raising the possibility that non-endocrine cells within the islet microenvironment contribute to the initiation of the proliferative response. Pericytes and endothelial cells are established sources of SCF (Ding et al., 2012), and islet endothelial cells in particular express high levels of SCF (Craig-Schapiro et al., 2025). As the number of endothelial cells increase, with the larger vessel diameter increases in obesity (Dai et al., 2013), endothelial SCF production could represent an important paracrine mechanism driving α-cell expansion.

More broadly, these findings raise the possibility that the primary cilium functions not only as a signaling platform for c-KIT but also as an important regulator of its mitogenic activity. This concept may have implications beyond pancreatic islets. Aberrant c-KIT signaling is a hallmark of several malignancies, including gastrointestinal stromal tumors, melanoma, mastocytosis, and subsets of acute myeloid leukemia (Abdellateif et al., 2023, Zhang et al., 2020). At the same time, the loss of primary cilia has emerged as a common feature of numerous cancers (Collinson and Tanos, 2025), including pancreatic ductal adenocarcinoma, where ciliary loss is thought to alter proliferative signaling pathways and promote tumor progression (Seeley et al., 2009, Kobayashi et al., 2017). Together, these observations raise the intriguing possibility that dynamic regulation of primary cilia may influence c-KIT-dependent proliferative responses in both normal physiology and disease. Determining whether ciliary remodeling contributes directly to c-KIT-driven tumorigenesis will be an important avenue for future investigation.

Collectively, our study provides a comprehensive ciliary proteome of αTC1-6 cells, identifies c-KIT as a previously unrecognized ciliary RTK, and demonstrate that c-KIT signaling depends on an intact primary cilium. In addition, our findings suggest that α-cell proliferation is regulated, at least in part, by paracrine signals originating from non-endocrine cells within the islet microenvironment and identify SCF/c-KIT signaling as a potential contributor to α-cell expansion and the development of hyperglucagonemia in obesity and type 2 diabetes.

## Author contributions

Conceptualization: JGK, LBP, STC. Data curation: GD and JGK. Formal analysis: GD, GDD, JGK. Funding acquisition: JGK, LBP. Investigation: GD, GDD, MGA, AK, TB, FC and GKP. Methodology: LBP, GD, and JGK. Resources: JGK, LBP, RBR, KB, RR, STC, LE. Supervision: JGK, LBP, RBR, KB, RR, STC, LE. Validation: LBP, JGK, RR, KB, STC, GD, LE. Visualization: JGK, GD. Writing Original Draft: LBP, STC and JGK. Writing – review and editing: LBP, JGK, LE, RR, KB, STC, GD, GDD, MGA, AK, TB and FC.

## Data availability

All data required to assess our conclusions are included in the manuscript. Raw images and source data for all figures can be obtained upon reasonable request to JGK.

## Acknowledgements

We thank Dorthe Nielsen for technical assistance, Professor Bernard Schermer from the Kidney Research Center Cologne, Germany, for kindly gifting the ARL13B-TurboID construct and Core Facility for Medical Proteomics, Medical Faculty University Tübingen for excellent assistance and support and the Quantitative Biology Center Tübingen (QBIC) for data management.

---

**Figure S1.**
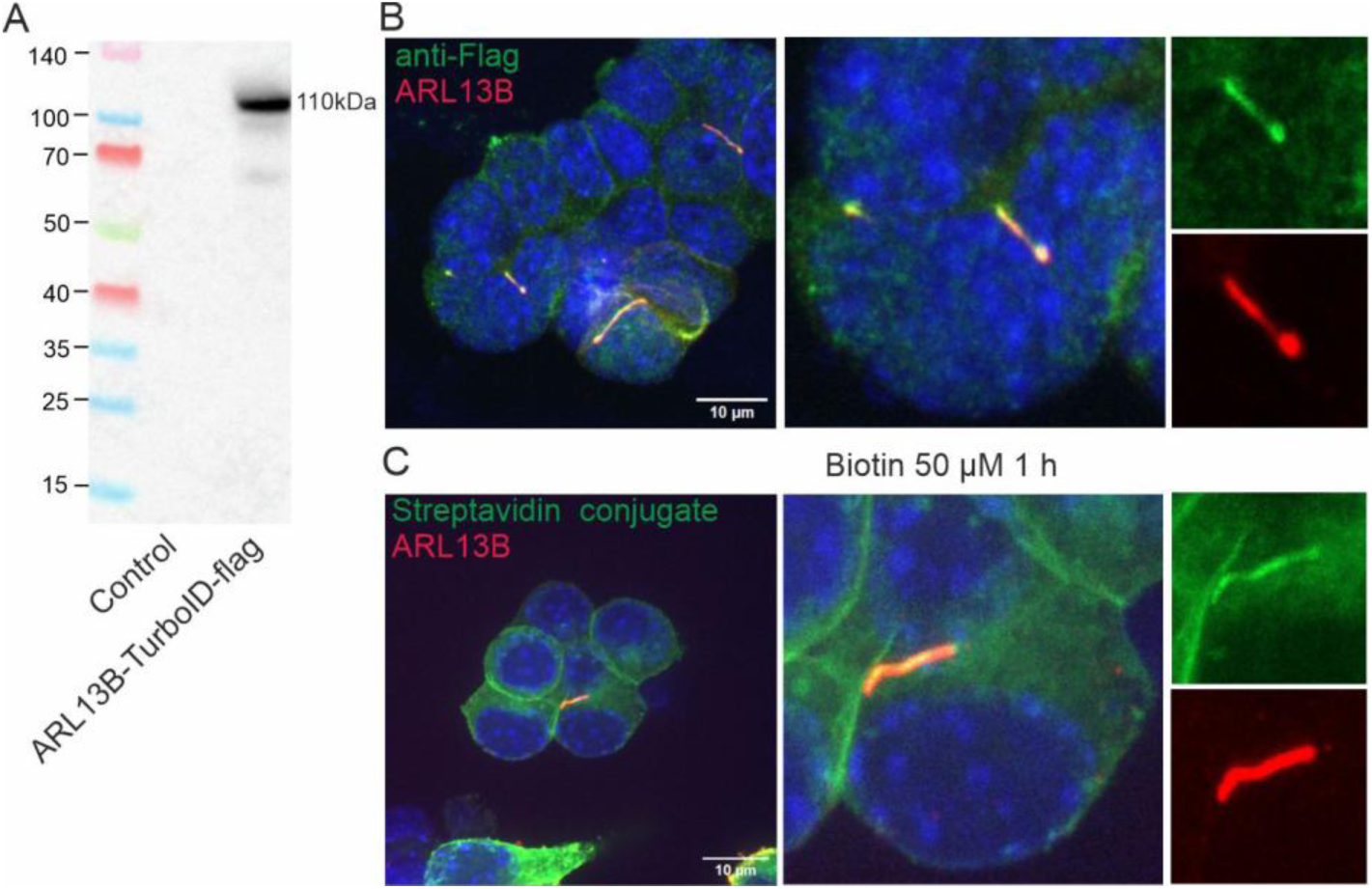
Confirmation of ARL13B-TurboID-3xFlag α-TC1-6 cell line function. (A) Representative western blot with control and α-TC1-6 cells, stably expressing ARL13B-TurboID-3xFlag. (B) Representative images of ARL13B-TurboID-3xFlag localization to primary cilia ARL13B (red), Flag (green). (C) α-TC1-6 cell stably expressing ARL13B-TurboID-3xFlag incubated for 1 h with 50 µM biotin showing Streptavidin staining of biotinylated proteins (green) and primary cilia stained with ARL13B (red).

**Figure S2.**
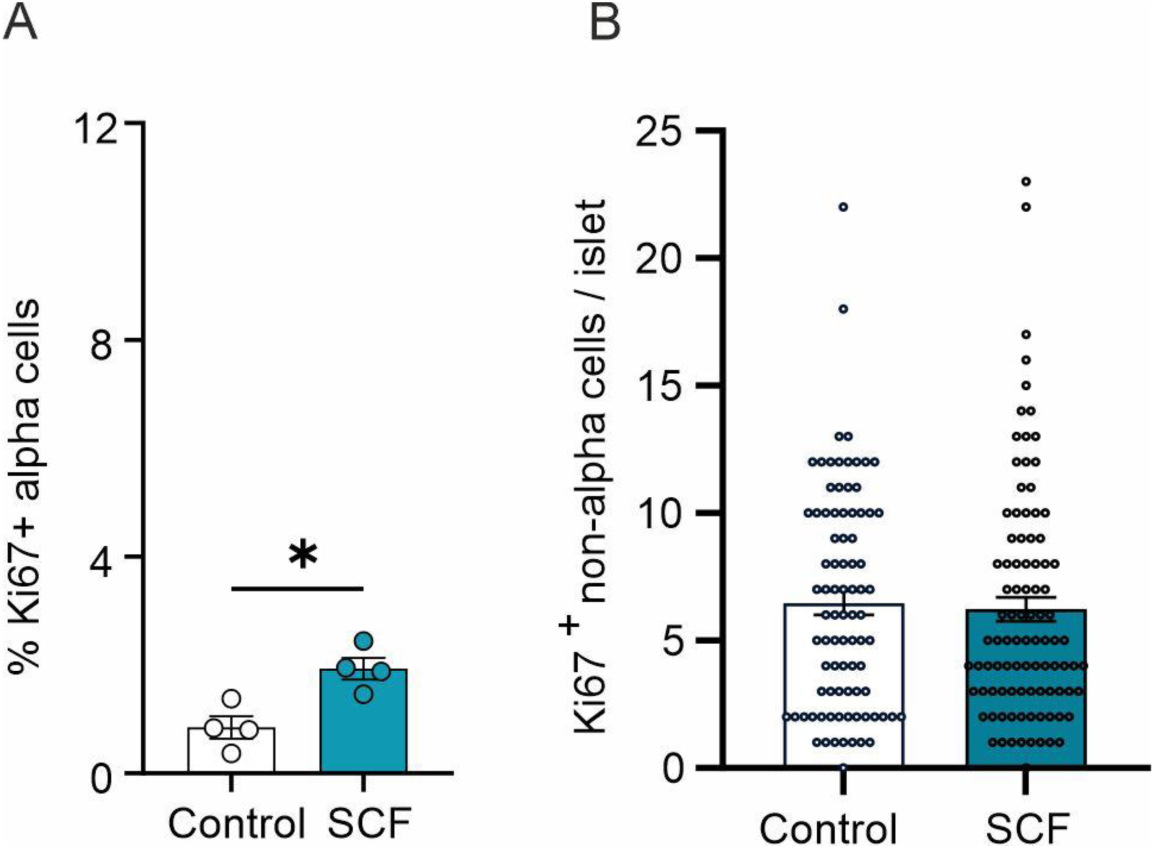
Stem cell factor expression in human islets correlates with BMI. Ki67 (A) alpha and (B) non-alpha cells in whole mouse islets treated with SCF for 72 or (n=4 islets from 3 mice). All data are presented as mean ± SEM, * (P<0.05), ** (P<0.01), *** (P<0.001).

**Figure S3.**
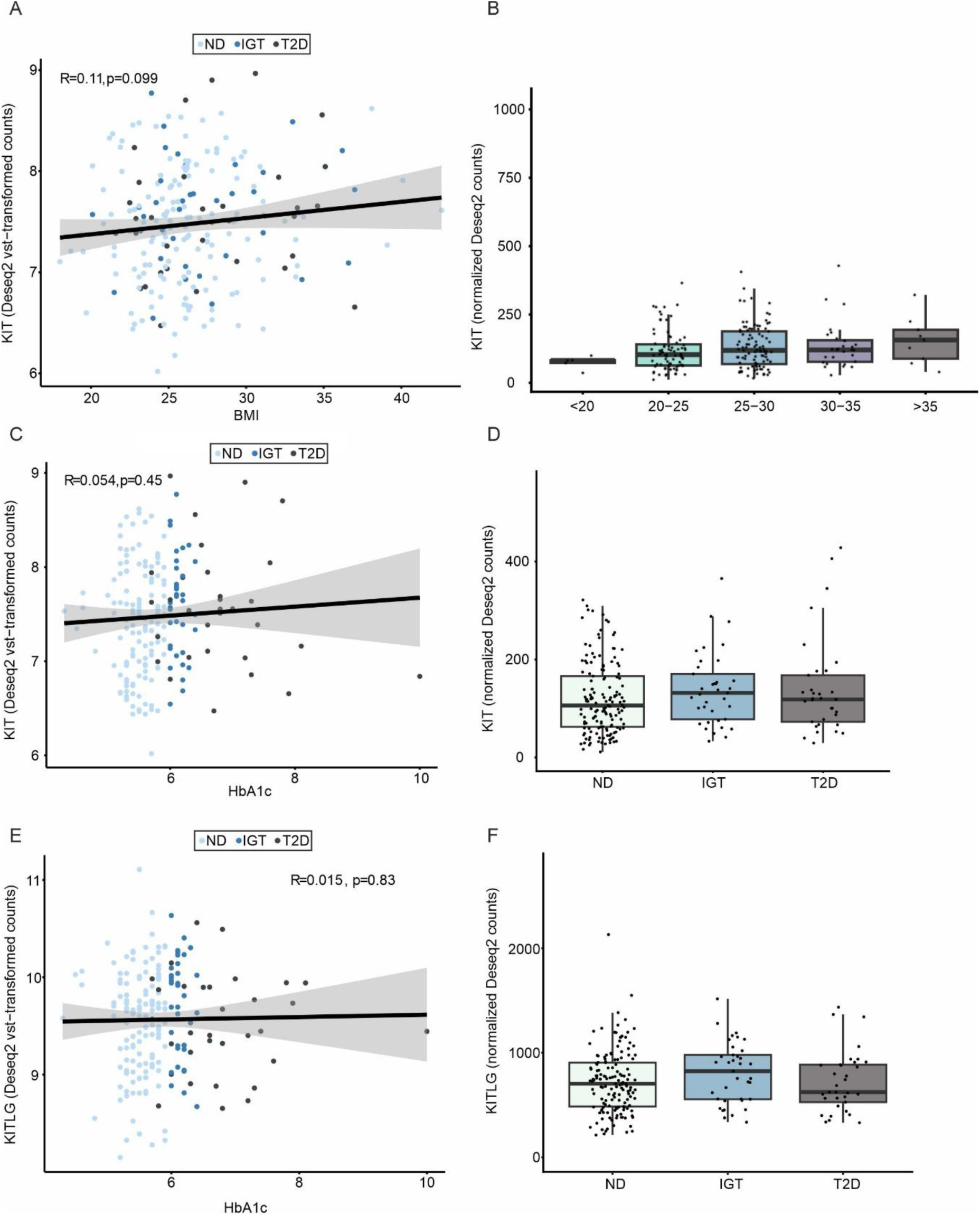
*KIT* and *KITLG* correlation with BMI and HbA1c. (A) Correlation between expression of *KIT* in whole human islets and body mass index (BMI) in non-diabetic subjects (ND), subjects with impaired glucose tolerance (IGT) or Type 2 diabetes (T2D). (B) Data form (A) stratified for BMI. (C) Correlation between expression of *KIT* in whole human islets and HbA1c in non-diabetic subjects (ND), subjects with impaired glucose tolerance (IGT) or Type 2 diabetes (T2D) D) Data form (C) stratified for in to ND, LGT and T2D. (E) Correlation between expression of *KITLG* in whole human islets and HbA1c in non-diabetic subjects (ND), subjects with impaired glucose tolerance (IGT) or Type 2 diabetes (T2D). (F) Data form (E) stratified into ND, LGT and T2D. All data are presented as Median ± 95%CI.

